# An automated method for precise axon reconstruction from recordings of high-density micro-electrode arrays

**DOI:** 10.1101/2021.06.12.448051

**Authors:** Alessio Paolo Buccino, Xinyue Yuan, Vishalini Emmenegger, Xiaohan Xue, Tobias Gänswein, Andreas Hierlemann

**Author notes:** These authors contributed equally to this work (Xinyue Yuan, Vishalini Emmenegger).

## Abstract

Neurons communicate with each other by sending action potentials through their axons. The velocity of axonal signal propagation describes how fast electrical action potentials can travel, and can be affected in a human brain by several pathologies, including multiple sclerosis, traumatic brain injury and channelopathies. High-density microelectrode arrays (HD-MEAs) provide unprecedented spatio-temporal resolution to extracellularly record neural electrical activity. The high density of the recording electrodes enables to image the activity of individual neurons down to subcellular resolution, which includes the propagation of axonal signals. However, axon reconstruction, to date, mainly relies on a manual approach to select the electrodes and channels that seemingly record the signals along a specific axon, while an automated approach to track multiple axonal branches in extracellular action-potential recordings is still missing.

In this article, we propose a fully automated approach to reconstruct axons from extracellular electrical-potential landscapes, so-called “electrical footprints” of neurons. After an initial electrode and channel selection, the proposed method first constructs a graph, based on the voltage signal amplitudes and latencies. Then, the graph is interrogated to extract possible axonal branches. Finally, the axonal branches are pruned and axonal action-potential propagation velocities are computed.

We first validate our method using simulated data from detailed reconstructions of neurons, showing that our approach is capable of accurately reconstructing axonal branches. We then apply the reconstruction algorithm to experimental recordings of HD-MEAs and show that it can be used to determine axonal morphologies and signal-propagation velocities at high throughput.

We introduce a fully automated method to reconstruct axonal branches and estimate axonal action-potential propagation velocities using HD-MEA recordings. Our method yields highly reliable and reproducible velocity estimations, which constitute an important electrophysiological feature of neuronal preparations.

## 1 Introduction

Axons are assumed to be faithful conductors of action potentials (APs) that encode and transmit information between individual neurons. Traditionally, axons are often considered as simple transmission cables, whose role is the reliable conveyance of APs to the presynaptic terminals of synaptically connected neurons [1]. Owing to recent technology advancements, such reductionist view of the role of the axon is being challenged. A growing body of evidence suggests that axons may provide important contributions to neuronal information processing [2, 3]. For example, the waveform of APs has been shown to be modulated during axonal conduction, which facilitated synaptic transmission to postsynaptic neurons [4]. Moreover, studies using two-photon imaging have found that structural changes of axonal arbors are involved in circuit-level mechanisms of perceptual learning [5]. Therefore, a precise tracking of axonal arbors, including the length of the axonal branches, number of branching points, and AP conduction velocities, will help to shed light onto mechanisms involved in axonal growth during development, axonal-AP modulation and their impact on neuronal signaling.

Due to the small diameters of axons of around 200 nm, the tracking of complete axonal arbors is challenging. Several classical electrophysiological techniques have been used for measurements and detection of AP propagation along axonal arbors. Whole-cell patch clamp, for example, has been used to measure the fidelity of AP propagation using dual patching at the soma and axonal blebs [6] or using cell-attached extracellular recordings in unmyelinated axons [7]. However, due to limitations in simultaneously recording from multiple sites along axons, the patch-clamp technique cannot be used to map axonal arbors. Alternatively, morphological information about neurons, including their axonal arbors, can be obtained with high-resolution imaging techniques. Recent advancements in imaging techniques, such as high-content imaging (HCI) [8, 9], have enhanced spatial resolution of acquired images. Together with the advances in image processing techniques [10, 11, 12], the reliability and throughput of such imaging methods allow for automatic tracing of neurites and their interconnections [12]. Yet, the use of imaging techniques requires fluorescent labels [13, 14], that may alter the physiological properties of the cells [15] through phototoxicity and photobleaching. In addition, it is difficult to extract axon morphologies in high-density cultures, where axons form bundles. HCI after post-hoc immunostaining ensures high spatial resolution, but axonal properties can only be investigated in live neurons.

High-density microelectrode arrays (HD-MEAs) have also been used to acquire electrophysiological signals of neurons at high temporal and spatial resolution [16]. Previous studies demonstrated the possibility to extract detailed representations of the extracellular electrical-potential landscape, so called “electrical footprints” of individual neurons from HD-MEA recordings by applying spike sorting and spike-triggered-averaging techniques [17, 18, 19]. These electrical footprints reflect the neurons’ morphology, so that researchers can use them for tracking neurite outgrowths of single neurons. However, the number of axonal arbors that could be extracted in the above-mentioned studies was limited to a few tens of cells in each sample due to tedious manual procedures to select and assign axonal signals. To date, no automatized method for extraction of morphological and functional information from large-scale electrophysiological HD-MEA data is available.

Building upon ideas and concepts of recent previous work [20, 21], we developed a novel, fully automated method to accurately reconstruct axonal arbors from functional electrophysiological HD-MEA recordings. Our method relies on a graph-based approach to reconstruct axonal branches and estimate AP conduction velocities. The proposed automatic method for reconstruction of axonal arbors and determining the corresponding AP conduction velocities from large-scale HD-MEA recordings opens up new possibilities to use axonal properties as electrophysiological biomarkers for studying compound efficacy and neural development as well as for drug screening and disease modeling.

## 2 Methods

In this section, we first introduce the biophysical simulation framework that we used as development test bench and for validation. Next, we describe in detail the implementation of the axonal tracking algorithm. Finally, we describe the protocols and procedures for experimental validation of our method.

### 2.1 Biophysical simulations

In order to develop and validate our axonal tracking approach, we initially used biophysical simulations. A simulation environment allowed us to explore complexities in the extracellular action potentials in a controlled manner, and to refine our method to deal with different cases. The simulations were carried out using `LFPy` 2.2.1 [22, 23] and `NEURON` 7.8.2 [24].

#### 2.1.1 Cell morphologies

We used morphological reconstructions of human pyramidal neurons from the Allen Institute of Brain Science cell-type database [25]. The cell models were downloaded from the Neuromorpho.org website [26] and included four samples (NeuroMorpho IDs: Cell 1 - [NMO_86990](http://neuromorpho.org/neuron_info.jsp?neuron_name=H16-03-006-01-04-03_563818992_m) - Figure 1A, Cell 2 - [NMO_86976](http://neuromorpho.org/neuron_info.jsp?neuron_name=H16-06-004-01-04-01_538906745_m) - Figure 1B, Cell 3 - [NMO_86965](http://neuromorpho.org/neuron_info.jsp?neuron_name=H16-03-006-01-04-03_563818992_m) - Figure 1C, Cell 4 - [NMO_87042](http://neuromorpho.org/neuron_info.jsp?neuron_name=H17-06-006-11-08-02_606834771_m) - Figure 1D). Since axonal tracking will be performed for cells cultured on a flat MEA substrate [19], the morphology of which extends principally in two dimensions, we modified the morphologies by setting all z-values to 0 µm, i.e., generated planar morphologies.

**Figure 1:**
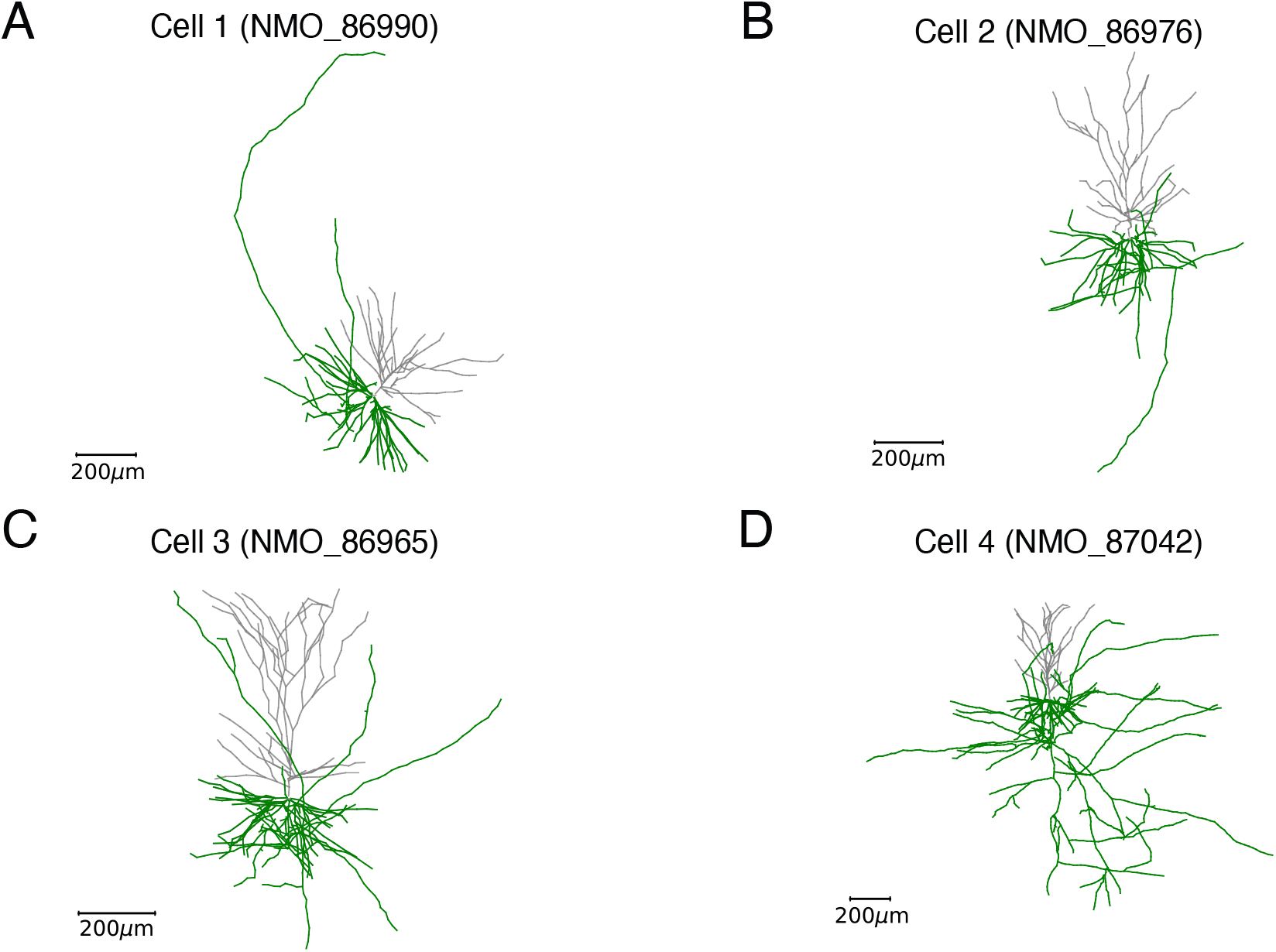
Neuron morphologies for biophysical simulations. Realistic morphologies from the Allen Institute of Brain Science cell-type database: Cell 1 (NMO_86990) - A, Cell 2 (NMO_860976) - B, Cell 3 (NMO_86965) - C, and Cell 4 (NMO_87042) - D. The axonal arbors are colored in green, while dendrites are in grey.

#### 2.1.2 Cell biophysics

For all cell models, biophysical properties were added in order to obtain realistic AP generation and axonal AP propagation. The membrane capacitance was set to 1 *µF/cm*^2^ for all compartments. Dendritic trees were defined to feature only passive membrane properties, with a membrane resistance of 150 *k*? and a reversal potential of −85 *mV*. The somatic compartments featured sodium and potassium Kv1 channels [27], with maximum conductances of 500 and 100 *S/cm*^2^, respectively. The axonal tracts also featured sodium- and potassium-channel conduction mechanisms, with maximum conductances of 500 and 400 *S/cm*^2^, respectively. The reversal potential for the sodium channel was set to 55 *mV*, and for the Kv1 channel to −98 *mV* [27]. *The axial resistance was set to 80 ?cm* and the temperature to 33° Celsius. The resting potential was set to −85 *mV*, and we simulated the cell model for 100 ms. The time step for the simulations was set to 0.03125 *ms*, yielding a sampling frequency of 32 *kHz*. In order to induce a single AP, we stimulated the cell with two to five consecutive synaptic inputs (`ExpSyn` mechanism - 1 ms between inputs) directly to the soma of the neuron.

#### 2.1.3 Modeling of extracellular signals

Extracellular potentials were modeled with a well-established forward-modeling scheme using the `LFPy` software [23]. Assuming a quasi-static, linear, isotropic, homogeneous, and infinite medium, the contribution of a neuronal transmembrane current *I*_*i*_(*t*), distributed over a line source (*line-source model*), centered at a point ***r***_*i*_ to the potential *ϕ*_*i*_(***r***_*j*_, *t*), measured by an electrode at position ***r***_*j*_, can be computed as:

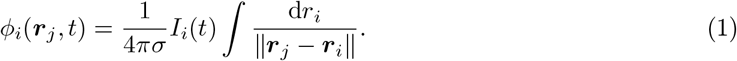

where *σ* is the extracellular conductivity (0.3 *S/m*). While the assumption of an infinite and homogeneous milieu is clearly violated in the presence of a highly insulating HD-MEA surface [28, 29], we did not apply any correction, e.g., by using the method of images [28]. A correction would only change the signal amplitudes but not alter signal timing and the relative signal amplitude distribution across the electrodes, which are pivotal for applying the proposed tracking algorithm.

The HD-MEA device was simulated using the `MEAutility` package [30], which is integrated in `LFPy` (version ≥2.1). A 100×100 electrode grid (10’000 electrodes in total) featuring a pitch of 17.5 µm, which represents the state-of-the-art of HD-MEA devices [31, 32, 33], was placed on the x-y plane at a vertical distance of 10 µm below the neuronal-cell plane. To represent the spatial extension of the electrodes, they were modeled as squares with a 5 µm side length. The *recorded* electrical potential was computed as the average over 10 points randomly positioned within the electrode surface using the so-called *disk-approximation* [22].

Figure 2A shows a visualization of the Cell 1 (NMO_86990) in black on top of the electrode grid of HD-MEA. Displayed is the complete morphology including axons and dendrites. The extracellular electrical-potential amplitude map (in log scale) is shown in Figure 2B, while the insets of Figure 2C, display the aligned intracellular and extracellular signals, showing the axonal propagation from the proximal part of the longest axon (blue, bottom) to the distal end (red, top).

**Figure 2:**
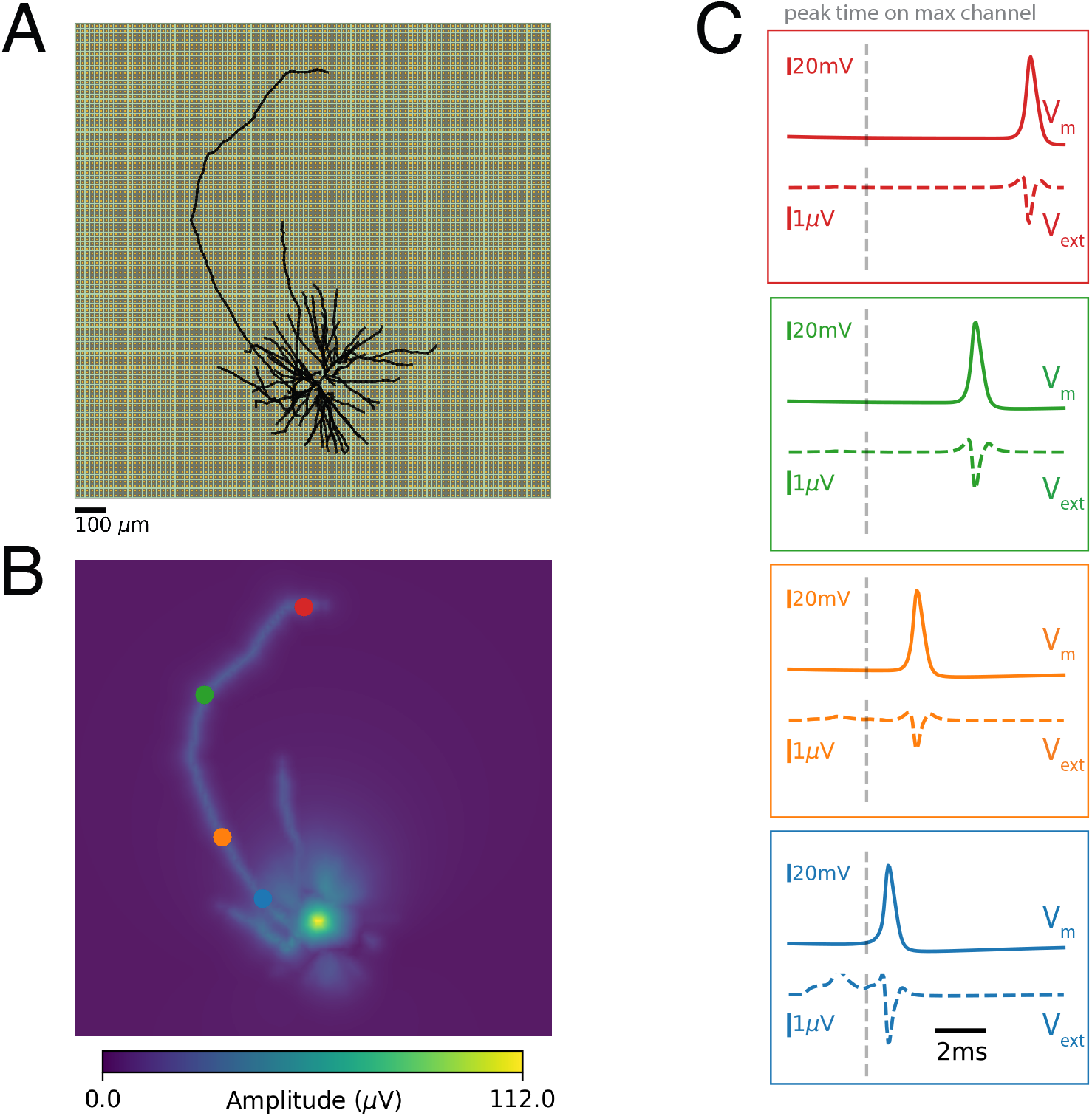
Simulation of extracellular signals. **A)** Representation of the HD-MEA and the *Cell 1* neuron located on top of the MEA. **B)** Amplitude map (in log scale) of the extracellular action potentials. Several axonal branches are clearly visible. **C)** Membrane potentials (*V*_*m*_ - top) and extracellular signals (*V*_*ext*_ - bottom) for the four points, indicated in color, along the longest axon in panel B. The vertical grey dotted line indicates the time of occurrence of the signal peak on the electrode featuring the largest AP amplitude.

### 2.2 Graph-based algorithm

*In this section, we describe the proposed algorithm, the application of which includes four main steps: i)* channel selection, *ii)* graph construction, *iii)* axonal-branch reconstruction, and *iv)* axonal-arbor pruning and velocity estimation. The method originated from ideas and concepts in our group [20, 34], which have been organized, modified, validated and assembled to obtain a coherent and fully functional method for axonal-arbor reconstruction. In Section 4 we compare the presented new approach to the previously used approaches.

#### 2.2.1 Channel selection

In order to track axonal branches, first, a subset of electrodes/channels needs to be selected that can be used for axonal tracking. Four *filters*, based on signal amplitudes, kurtosis, peak time standard deviations, and initial signal delays are available. An appropriate channel selection depends on many factors, such as the probe geometry and the noise level, therefore, the proposed method gives freedom to the user to modify the filter configuration to maximize tracking performance. In the following section, we briefly describe how the different filters operate and we display, in Figure 3, the channel selection for a real neuronal footprint, which was obtained in an HD-MEA recording using ∼20’000 channels [20]). The footprint is shown in Figure 3A, the channel selection for each available filter in Figure 3B-E and the selection using all 4 filters in Figure 3F.

**Figure 3:**
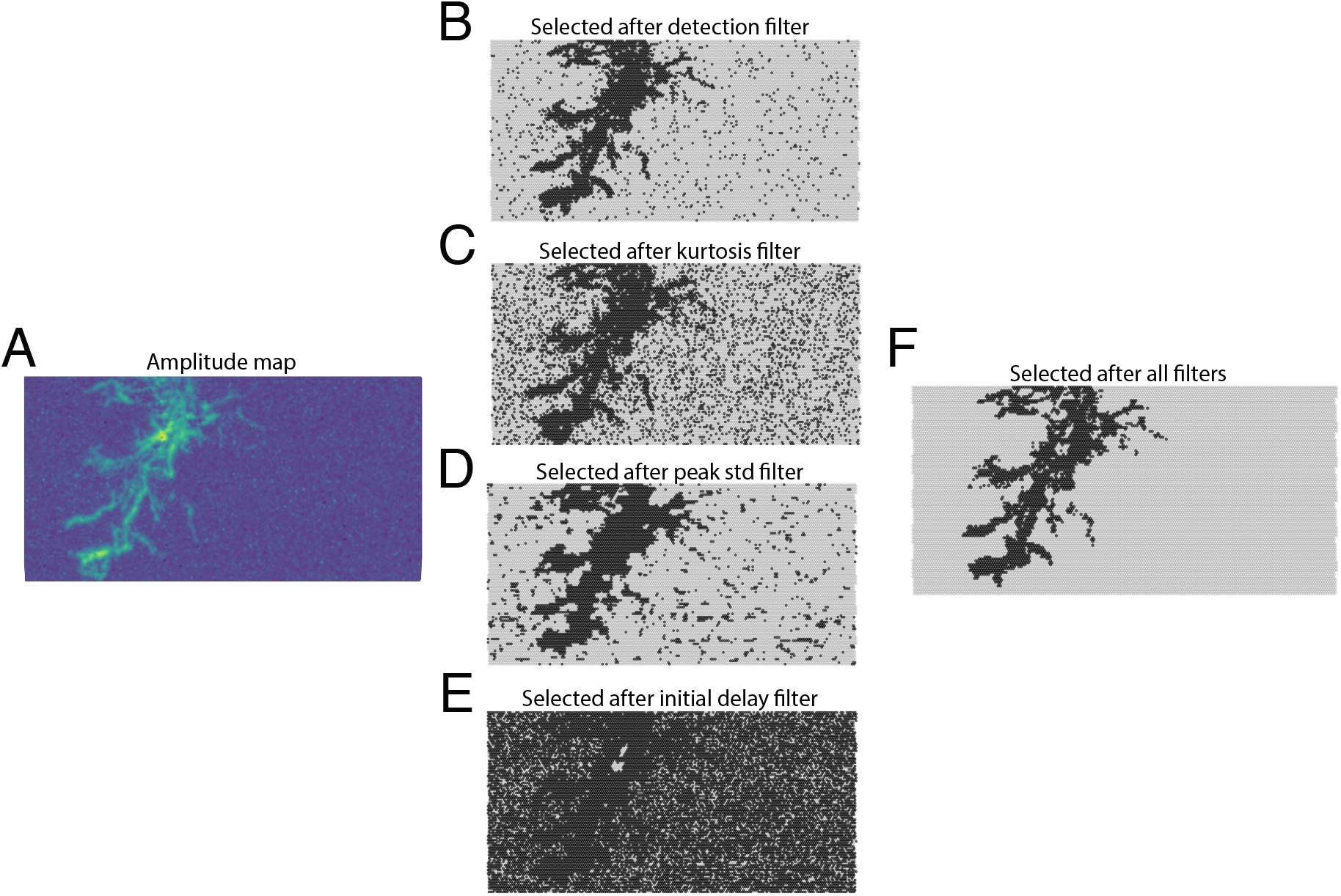
Channel selection procedure. **A)** Amplitude map (log scale) of a neuron footprint. **B-E)** Selected (black) and excluded (grey) channels after filtering for amplitude (B), kurtosis (C), peak time standard deviation (D) and initial delay (E). Selected (black) channels after combining all filters.

##### Amplitude filter

The first available filter is a *detection filter* based on the amplitude of the recorded signal. Only channels with a peak-to-peak amplitude larger than a detection threshold are kept for further processing. The detection threshold can be defined relative to the largest signal amplitude of the recording (default) or as an absolute value in µV, and the default setting is 0.01 (1 %). Figure 3B shows all available channels in grey and the selected channels after applying the detection filter in black.

##### Kurtosis filter

Second, a filter based on kurtosis can be used in order to ignore channels that may contain only noise. A noisy channel, in fact, may pass the detection filter unnoticed. However, if a channel features *signal spikes*, its kurtosis should be above zero, i.e., it should exhibit a supergaussian distribution. The default setting of the kurtosis filter is 0.3, and all channels with a kurtosis value below this threshold are removed. Figure 3C shows all available channels in grey, and the selected channels after application of the kurtosis filter in black.

##### Peak time standard deviation filter

A third available filter relies on the standard deviation of the occurrence time of the signal peaks in neighboring channels (channels within 30 µm distance are selected as default) [18]. The recommended threshold for this filter is 1 ms, and the channel selection based on this filter is shown in Figure 3D.

##### Initial delay filter

Finally, since our aim is to track axons, we remove all channels whose the signal peak occurrence time is lower than that of the channel featuring the largest signal amplitude, referred to as *initial channel* plus an additional delay (set to 0.1 ms by default). The signal in this *initial channel* is assumed to originate from the axon initial segment [17]). This removal is done to *wait* until electrical-signal propagation has entered the axonal branches. Figure 3E shows all available channels in grey, and the selected channels after using the initial-delay filter in black.

All selection filters are applied separately, and the final channels selected correspond to the intersection of the channels selected by each individual filter (Figure 3F). Finally, isolated channels (selected channels without a *neighbor* within 100 µm distance) are removed from the selection.

#### 2.2.2 Construction of the graph

After the channel selection, the *remaining* channels are used as the nodes of a graph. Prior to the graph construction, however, the channels are sorted based on the following heuristic:

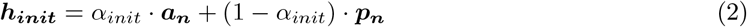

where ***a***_***n***_ are the normalized amplitude values, ***p***_***n***_ are the normalized peak latencies, and *α*_*init*_ is a scalar that weighs the contributions of amplitude and peak latency. The channel sorting will influence the order of initial channels chosen to construct axonal branches. By default, *α*_*init*_ is set to 0.2, so that the channels with signal-amplitude peaks that occur comparably late are favored, and among those, the channels featuring the largest amplitudes.

When channels are sorted, they are used as *nodes* to populate a directed graph. The graph is built using the `NetworkX` Python package [35]. Next, *edges* are added to the graph. For each node, at most `n_neighbors` edges (default: 3) can connect to other candidate nodes if: *i)* the candidate node has a signal peak occurring earlier in time, and *ii)* the candidate node is within a maximum distance (default: 100 µm). Among the candidate nodes that satisfy these two requirements (there can be more than `n_neighbors` depending on the electrode density of the MEA), the channels featuring the largest amplitudes and the lowest distances are favored. Channels for which there is no other channel with an earlier peak occurrence (excluding the initial channel) are connected to the initial channel if they are within a defined spatial range (default: 200 µm distance). Each edge is added to the graph along with the average amplitude of the nodes that it connects (edge amplitude). After all edges have been added to the graph, all amplitudes values are retrieved and normalized between 0 and 1. For amplitudes, the slope is reversed so that the largest amplitude has a value of 0, and the smallest one is assigned a value of 1. We denote these normalized edge amplitudes as ***h***_***edge***_, since they are used as a heuristic to find axonal branches. For all edges connecting to the initial channel, the *h*_*edge*_ value is set to 2.

The graph nodes for the model of *Cell 1* is shown in Figure 4A. The nodes are colored according to ***h***_***init***_ values. Figure 4B shows the edges colored according to ***h***_***edge***_ values for the same cell model. It becomes evident that the ***h***_***init***_ values exhibit local maxima at the axon ends and that lower values of ***h***_***edge***_ nicely coincide with axonal paths. These two heuristics are then used to reconstruct the axonal branches.

**Figure 4:**
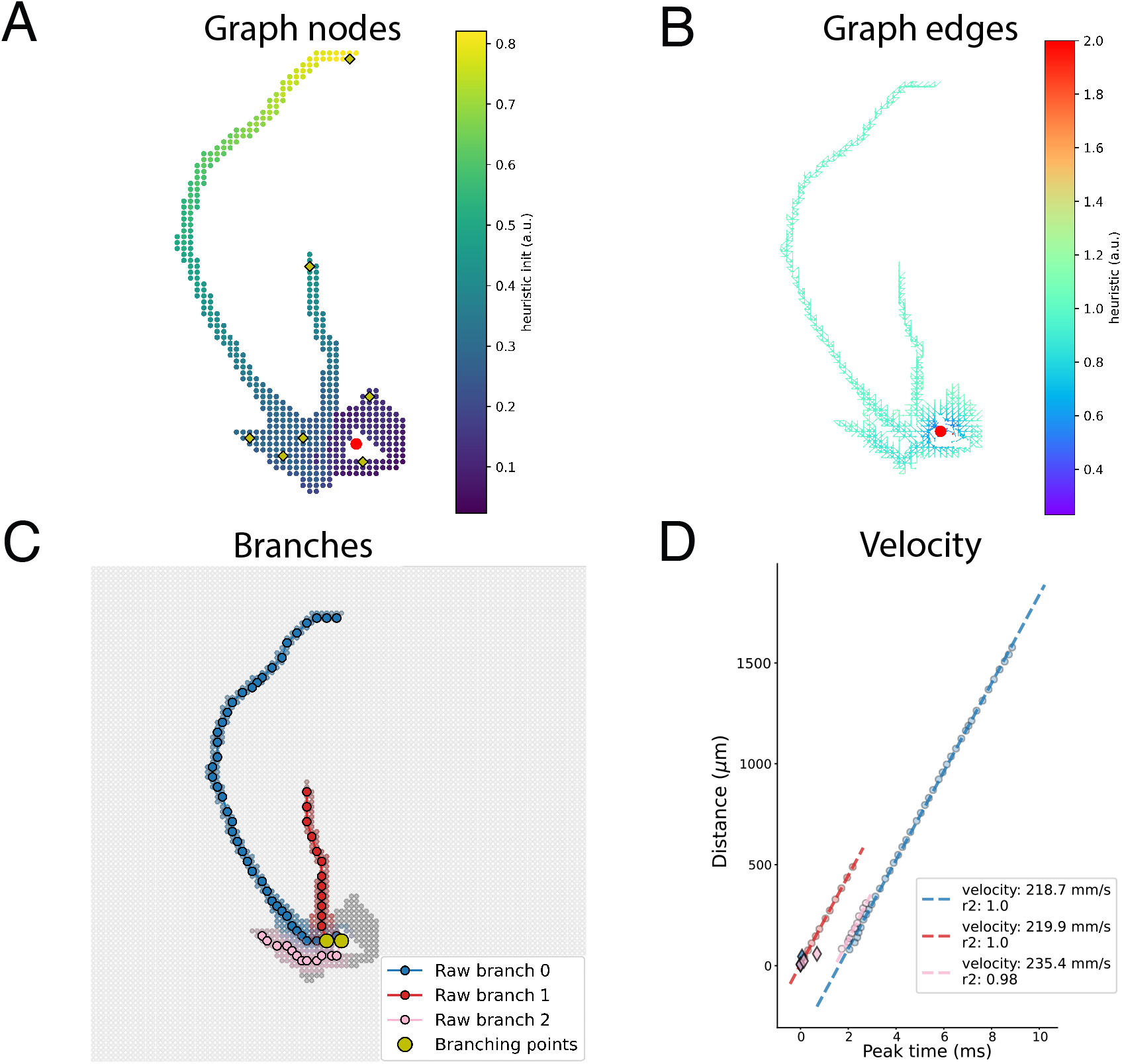
Axonal reconstruction method. **A)** Graph nodes colored according to ***h***_***init***_ values for the *Cell 1* neuron. The nodes marked with a yellow diamond indicate nodes for which a path towards the initial channel has been searched for. **B)** Graph edges colored according to ***h***_***edge***_ values. **C)** Identified raw axonal paths. The dark grey dots are the selected channels. The colored nodes around an identified path are the neighbor nodes for that path, which have been removed for further searches. The yellow circles indicate the branching points. **D)** Robust velocity estimation. For each reconstructed branch, a robust estimator was used to fit the axonal AP propagation velocity. The blue and pink diamonds at the bottom left show detected outliers from branch 0 (blue) and branch 2 (pink) respectively, which were removed from the cleaned paths.

#### 2.2.3 Axonal branch reconstruction

The two heuristic functions (***h***_***init***_ and ***h***_***edge***_) are used to reconstruct axonal branches. The goal of this step is to find possible paths that represent axonal branches. Since graph nodes are already sorted by *h*_*init*_, this procedure loops through the nodes and attempts to find paths ***P*** towards the initial channel while minimizing the edge heuristic ***h***_***edge***_. A path is searched between a node and the initial channel only if the node is a local maximum in the ***h***_***init***_ space, i.e., it has the largest value of *h*_*init*_ compared to other channels within a distance of 100 µm (distance adjustable by the user). This approach ensures that only a small number of paths is reconstructed and improves the efficiency of the method. The nodes indicated in red in Figure 4A represent the local maxima that have been identified as starting nodes for axonal branches. The shortest path is obtained using the *A*\* method, which also considers the spatial distance between channels. In order to avoid *long jumps*, the distance used by the algorithm is computed as:

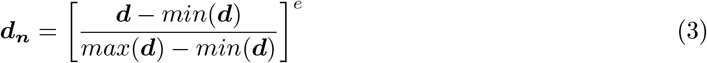

where *d* is the distance between two nodes that are connected by an edge and *e* is the configurable exponential (2 by default). To further minimize *long jumps*, the value of *e* can be increased. Finally, the *A*\* method finds the path that minimizes:

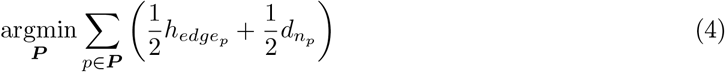

where *p* is a single node, and ***P*** is the set of nodes that makes up a path. When a path is found, the channels within the neighborhood of each channel in the path (by default, within 100 µm) are stored as the *neighbor channels* set. If a channel in a newly identified path is already included in the set of neighbor channels (i.e., it is neighboring an already existing path), this channel and all channels in the path closer to the *initial channel* are removed from the path. The last channel, which has not been removed, is connected to the closest node of the already identified closest path. In this case, the new channel becomes a *branching point*. After all paths and all branching points have been found, paths are pruned and merged. A path is pruned if a portion of it extending from a branching point does not have at least three points by default (this value is adjustable by the user). Finally, pairs of paths that, after pruning, share a branching point which corresponds to the last channel of one path and the first channel of the other path are merged. After pruning and merging, a path is stored as a *raw* axonal branch if two conditions are met: *i)* the length of the path is larger than a path length threshold (default is 100 µm), and *ii)* the path contains at least a minimum number of points (5 by default). Once a path has been accepted, all the channels of the path and the ones within a *neighbor radius* (50 µm by default) of any of its nodes are stored in the memory and excluded from further searches. This step ensures that no duplicate paths are found for the same axonal branch.

The identified branches for the *Cell 1* model are shown in Figure 4C. In this case, three raw branches were found (blue, red, pink). The grey dots are the selected channels and the shaded nodes around each path (with the same color) indicate the channel neighbors, which were removed from further path searches. The yellow circles represent the branching points.

The full algorithm to estimate raw branches is described in Algorithm 1 in Appendix A.

#### 2.2.4 Path cleaning and velocity estimation

After obtaining the set of paths, axonal velocities can be estimated. Peak times are computed as the difference between the occurrence time of the signal peak at each node and the peak time occurrence of the first node in the path (which is the one featuring the earliest signal peak by definition). Cumulative distances are calculated by integrating the distances between the channels along the path.

Once peak times and distances have been computed, a robust linear fit using the Theil-Sen regressor (using `scikit-learn` [36]) is used to reduce contributions of possible outliers. The velocity estimate is derived from the slope of the regression line. We use a non-parametric and robust approach to identify and remove possible outliers from the path. We first compute the prediction error for each channel. We then identify outliers as nodes with an error of *N* times larger than the median absolute deviation (MAD) of the error distribution (*N* is 8 by default) and is above a fixed threshold (30 µm by default). Outliers are then removed from the axonal branches, and a new linear fit is computed. In some cases, it could happen that a path presents a *shortcut* either between different branches or within the same branch with an undetected axonal section. In this case, a jump in the peak signal occurrence times is observed. In order to correct for this unwanted behavior, the method attempts to split the path to fit the sub-paths separately, when jumps in the peak times are detected (>1 ms by default). If the average *R*^2^ of the sub-paths is larger than the *R*^2^ of the original path, the path is split and the sub-paths are considered as separate branches. Finally, axonal branches with an *R*^2^ value below a user-defined threshold (default 0.9) are discarded.

Figure 4D shows the peak latencies (x-axis), cumulative distances (y-axis), and the fitted AP propagation velocities (dashed lines) for the raw branches displayed in Figure 4C for the *Cell 1* model. The linear fit achieves a very high *R*^2^ value, partially owing to the removal of outliers of the blue and pink branches, depicted as diamond shapes.

### 2.3 Software implementation and code availability

The implementation of the above-described algorithm is available as an open-source Python package called `axon_velocity` on GitHub (https://github.com/alejoe91/axon_velocity) and on PyPi (https://pypi.org/project/axon-velocity/). All the code needed to reproduce figures in this article can be found in the figure_notebooks folder of the GitHub repo, while the required data are available at Zenodo (https://doi.org/10.5281/zenodo.4896745).

The graph-based algorithm takes the electrode array template (a `numpy` array with dimensions num_channels x num_samples), the x-y electrode locations (a `numpy` array with dimensions num_channels x 2), and the sampling frequency as required arguments. Additionally, all algorithm-specific parameters can be passed as extra arguments:

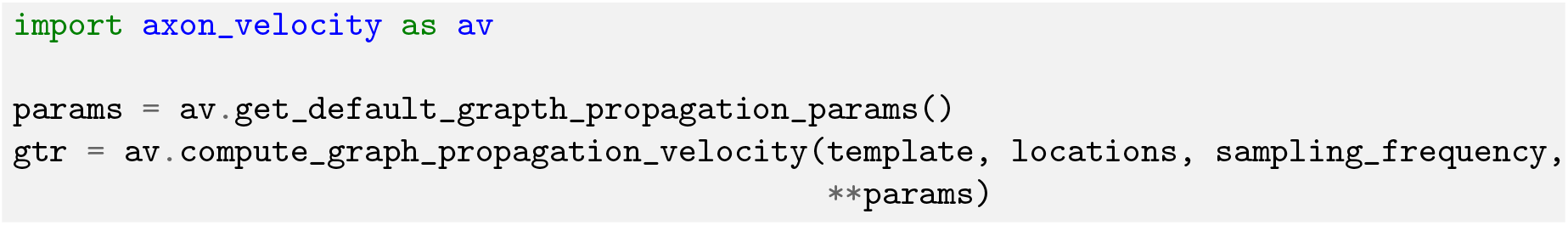

The returned `gtr` object is a `GraphAxonTracing` object, which contains the following fields:

- `branches`: list of dictionaries for the detected axonal branches. Each dictionary contains the following fields.
  - `channels`: selected channels in the path
  - `velocity`: velocity estimate in *mm/s*
  - `offset`: offset (intercept) of velocity estimate
  - `r2`: *r*^2^ of the AP velocity fit
  - `error`: standard error of the linear fit
  - `pval`: p-value of the linear fit
  - `distances`: array with cumulative distances computed along the branch
  - `peak_times`: array with signal peak occurrence time differences to initial channel

- `selected_channels`: list of selected channels used for axonal tracking
- `graph`: the `NetworkX` graph used to find axonal branches

In Table 2 of Appendix B we list and describe the additional parameters (`**params`), their default values, their types, and a brief description of their role.

### 2.4 Experimental procedures

#### High-density microelectrode arrays

To validate the tracking algorithm with experimental recordings, we used data from two types of HD-MEA chips: the first device features 26’400 electrodes with a center-to-center electrode distance of 17.5 µm and can record from up to 1024 channels simultaneously at 20 kHz [31, 32] (referred to as MEA1k); the second device is a dual-mode HD-MEA including switch-matrix and active-pixel readout schemes for electrodes [37, 20] (referred to as DualMode). It features a full-frame readout of 19,594 electrodes at a sampling rate of 11.6 kHz; the center-to-center electrode distance is 18 µm.

#### Cell cultures and plating

Rat primary neurons were obtained from dissociated cortices of Wistar rats at embryonic day 18, using the protocol described in Ronchi et al. [21]. All animal experimental protocols were approved by the Basel-Stadt veterinary office according to Swiss federal laws on animal welfare and were carried out in accordance with the approved guidelines.

Prior to cell plating, HD-MEA chips were sterilized using 70% ethanol for 30 minutes. Ethanol was then removed, and the chips were rinsed three times with sterile tissue-culture-grade water. The HD-MEA chips were coated with a layer of 0.05% polyethylenimine (Sigma) in borate buffer to render the surface more hydrophilic. On the plating day, a layer of laminin (Sigma, 0.02 mg/mL) in Neurobasal medium (Thermo Fisher Scientific) was added on the array and incubated for 30 minutes at 37 °C to promote cell adhesion. We dissociated cortices of E-18 Wistar rat enzymatically in trypsin with 0.25% EDTA (Gibco), followed by trituration. Cell suspensions of 15,000 to 20,000 cells in 7 *µ*L were then seeded on top of the electrode arrays. The plated chips were incubated at 37 °C for 30 min before adding 2 mL of plating medium. The plating medium consisted of Neurobasal, supplemented with 10% horse serum (HyClone, Thermo Fisher Scientific), 0.5 mM Glutamax (Invitrogen), and 2% B-27 (Invitrogen). After 3 days, 50% of the plating medium were replaced by a growth medium, which consisted of D-MEM (Invitrogen), supplemented with 10% horse serum, 2% B27, and 0.5 mM Glutamax. The procedure was repeated twice a week. The chips were kept inside an incubator at 37°C and 5% CO2. All experiments were conducted between days in vitro (DIVs) 10 and 28.

#### Extracellular recordings and analysis

For the MEA1k system, only 1024 channels of the array’s 26’400 electrodes can be recorded simultaneously, therefore an *axon scan* assay was performed: we sequentially recorded 33 different configurations of randomly placed electrodes in order to cover the entire chip area, while the 200 electrodes showing the highest spontaneous activity were fixed. Each configuration was recorded for 120 seconds. The recorded data were analyzed using SpikeInterface [38]: the signals from the fixed electrodes were concatenated in time and spike-sorted using Kilosort2 [39]. The spike sorting output was automatically curated by removing units with a firing rate lower than 0.1 Hz, an ISI violation threshold [40] higher than 0.3, and a signal-to-noise ratio lower than 5. Afterwards, the automatically curated data was exported to Phy [41, 42] for visual inspection and manual curation. The manually curated data were then used to extract full templates across the entire array: first, the spike trains were categorized depending on the start and end time of the different configurations; second, the template for each configuration was computed as the median of all extracted waveforms; finally, templates extracted from different configurations were averaged to obtain the final full template.

For the DualMode system, we analyzed a short full-frame recording of ∼285 seconds. As most spike sorters do not handle more than ~ 1000 channels, we first computed the spike rate of each channel using a spike detection based on 5 times the median absolute deviation. We then selected the 1024 most active channels and spike sorted them using Kilosort2 and the same automatic curation as for the MEA1k recordings. A final manual curation step using Phy was performed, and templates were extracted by combining spike times and the full-frame recording.

### 2.5 Evaluation of the tracking performance

In order to evaluate the performance of the proposed axon-tracking algorithm, we used the simulated data as ground truth. The ground-truth branches of the cell models were matched to the estimated axonal branches using a many-to-one strategy (since the estimated branch could span over one or more ground-truth branches). The matching was performed by computing the median distance of each ground-truth path to each estimated path. A possible match was called if the median distance was below a 40 µm threshold. Among the ground-truth branches matched to the same estimated branch, overlapping ground-truth branches were discarded. Overlapping branches were defined as ground-truth branches with more than 20% of their segments being located within a distance of 15 µm. In case overlapping ground-truth branches were found, the shortest ones were removed.

After the matching procedure, tracking errors and AP propagation velocities were computed for each estimated branch. The tracking errors were computed as the distance between each channel of an estimated branch and the closest segment of the matched ground-truth branches. Tracking errors were reported as mean±standard deviation in Table 1. In case of velocities, we also computed the absolute velocity error (*abs*(*v*_*gt*_ − *v*_*est*_)) and the relative velocity error (*abs*(*v*_*gt*_−*v*_*est*_)*/v*_*gt*_). Here *v*_*gt*_ is the ground-truth velocity – computed as the weighted average of the branch AP propagation velocity with respect to the branch length – and *v*_*est*_ is the estimated branch AP propagation velocity.

**Table 1:**
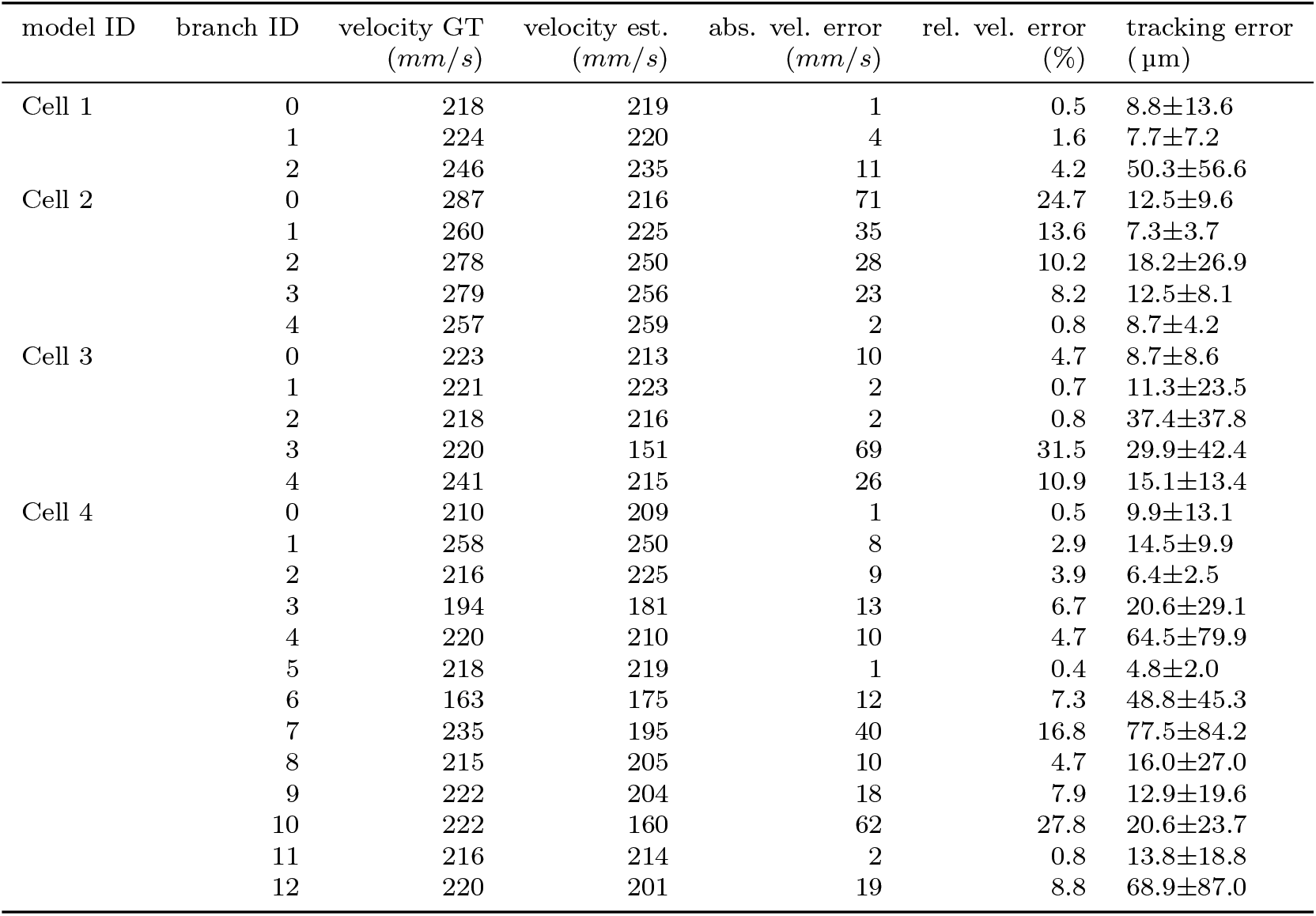
Performance on simulated data of model cells. Each entry of the table reports the Cell model (1, 2, 3, 4), the branch IDs (corresponding to Figure 5), the ground-truth and estimated velocities, the absolute velocity error (in *mm/s*) and the relative error (in %). The last column displays the mean and standard deviation of the tracking error in µm.

## 3 Results

### 3.1 Algorithm performance on realistic, simulated morphologies

In order to validate and assess the performance of the proposed method, we analyzed the axonal reconstructions and velocity estimations of simulated extracellular APs using the realistic morphologies from the Allen Institute database (Figure 1). Already from the morphologies, one can appreciate that the first three neuronal models (Cell 1, Cell 2, Cell 3) displayed well separated axonal branches, while Cell 4 (Figure 1D) showed a much more intricate axonal arborization.

We ran the graph-based algorithm with default parameters (listed in Table 2) and evaluated the tracking results against ground-truth information of the model cells. Figure 5 shows the estimated branches as dots and the matched ground-truth branches as lines. The estimated and corresponding matched ground-truth branches are plotted in the same color. Qualitatively, the developed method correctly identifies the main axonal branches of all tested model cells and shows good performance even for Cell 4, despite the multitude of axonal branches crossing each other. Table 1 shows the ground-truth and estimated velocity, the absolute and relative velocity errors, and the mean and standard deviation of the tracking errors for all estimated branches of the four model cells. In most cases (19 out of 26 axonal branches) the relative error is below 10 %. Higher velocity and tracking errors can be due to a partial match to the ground-truth branch (e.g., branch 0 in Figure 5B and branch 3 in Figure 5C). Nevertheless, the proposed tracking algorithm is capable of correctly reconstructing large portions of the axonal arborization of all model cells.

**Figure 5:**
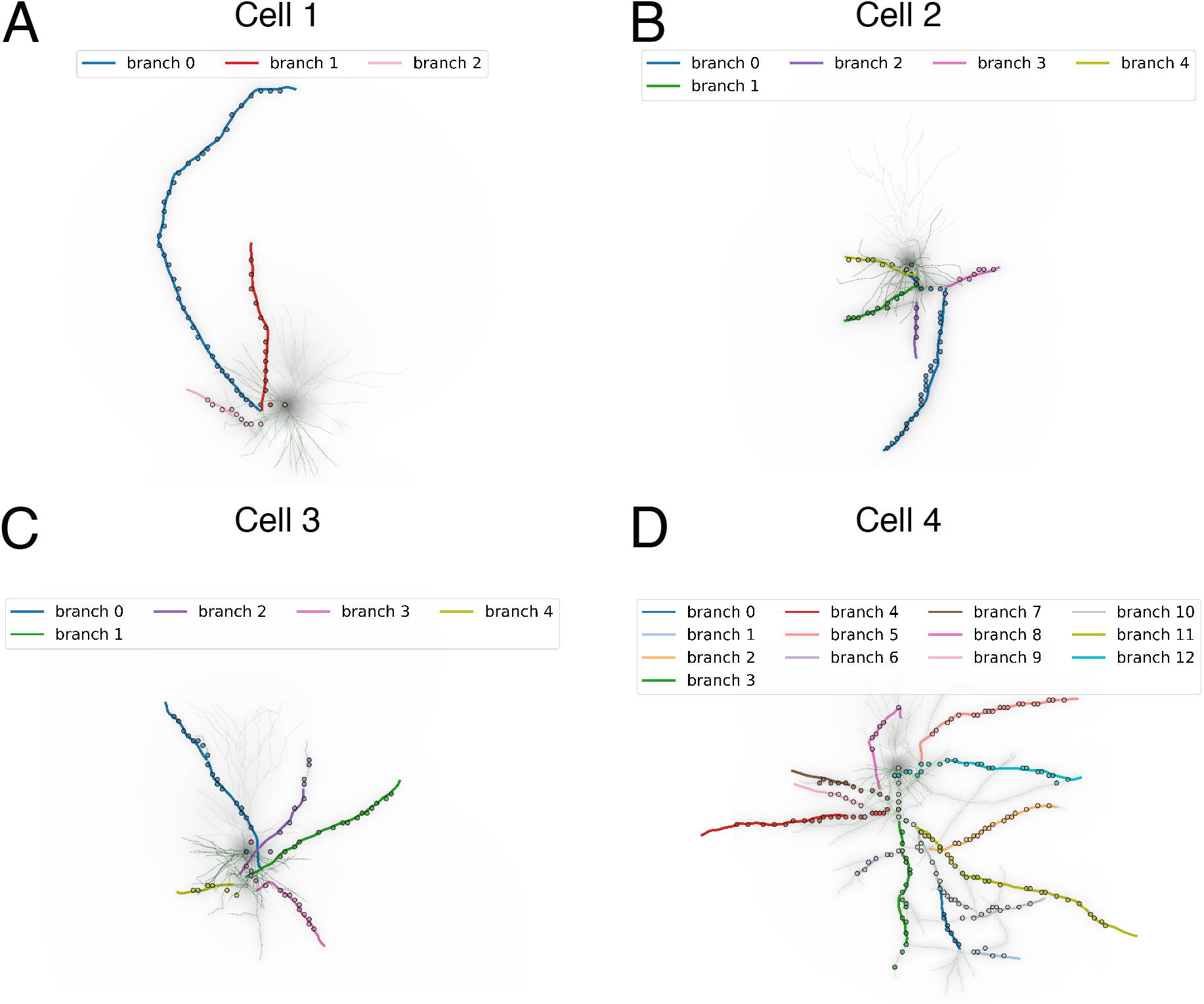
Axonal reconstruction on realistic neuron morphologies. Morphological reconstructions of Cell 1 **A)**, Cell 2 **B)**, Cell3 **C)** and Cell 4 **D)**. Colored lines display ground-truth branches that have been matched to the reconstructed branches (colored circles). The morphology of the cell is shown in the background.

### 3.2 Application to HD-MEA recordings

After validating the tracking performance of the proposed algorithm on simulated data, we analyzed experimental data from recording sessions, of two different HD-MEAs, a MEA1k and a DualMode recording. In both cases, we ran the proposed tracking algorithm using a detection threshold of 1%, a kurtosis threshold of 0.1, a standard deviation threshold of the signal peak occurrence time of 0.8 ms, and an initial delay of 0.2 ms. For the MEA1k dataset, the spike-sorting procedure after manual curation yielded 77 isolated units. Out of these, 67 units had detectable axonal branches. The algorithm found a total of 249 axonal branches, with velocities of 386.03 ± 250.7 *mm/s*, path lengths of 458.07±257.03 µm and *R*^2^ values of 0.94 ± 0.05. In Figure 6A we show all reconstructed axonal branches with a visualization of the MEA1k device with 26’400 electrodes in the background. Figure 6B shows a representative neuron of Figure 6A (marked in blue). The amplitude map of the template (top left), the peak latency map (top right), the reconstructed branches (bottom left), and the fitted velocities (bottom right) are shown. For this neuron, the channel selection yielded 1252 channels, featuring 8 axonal branches with path lengths of 486.77 ± 151.15 µm, AP propagation velocities of 417.57± 116.65 *mm/s*, peak-to-peak extracellular amplitude of 69 µV, and *R*^2^ values of 0.93±0.05.

**Figure 6:**
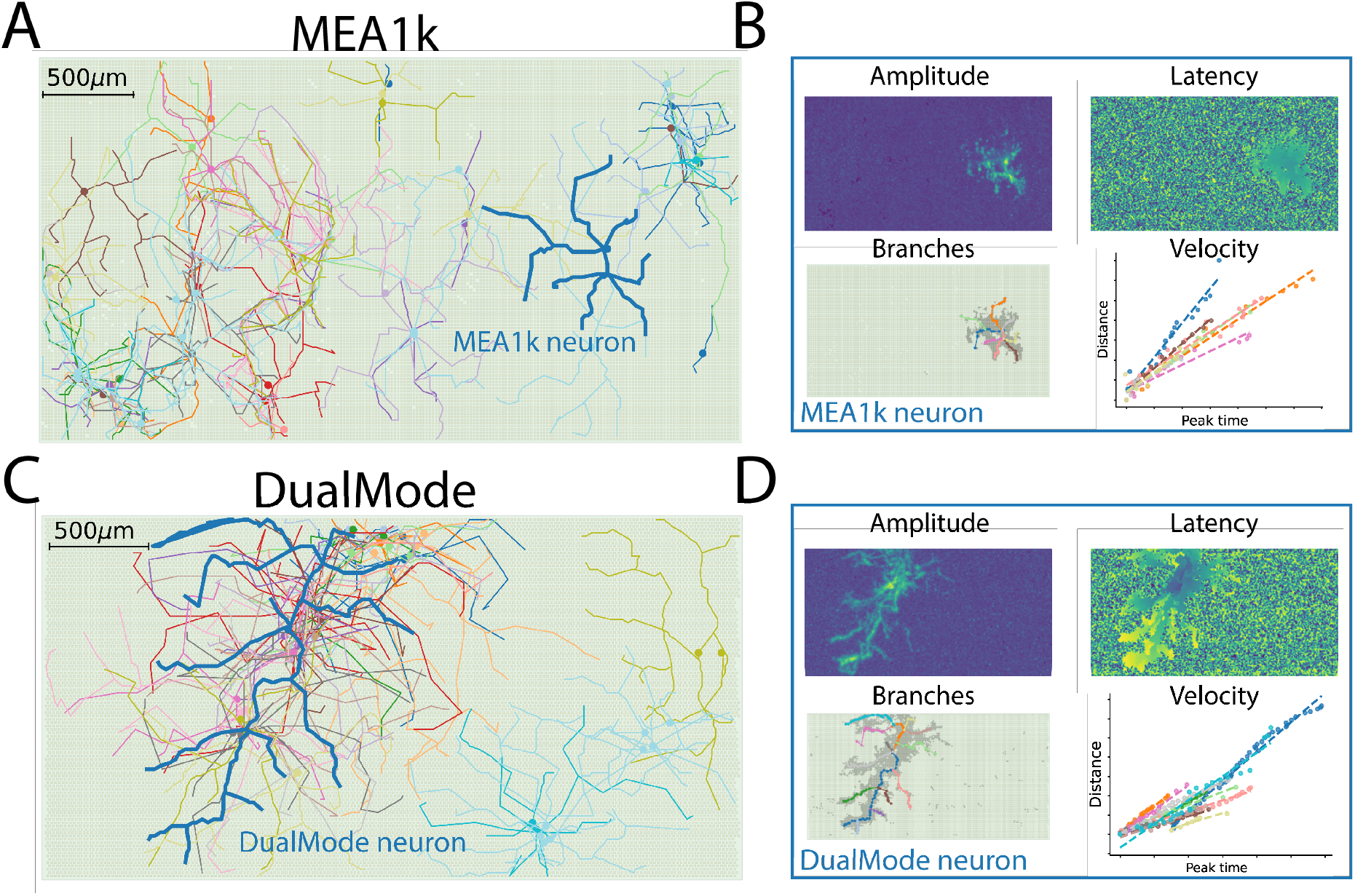
Application to HD-MEA recordings. **A)** Axonal arbors on a MEA1k device. All reconstructed units are displayed with a representation of the MEA (26’400 channels) in the background. **B)** Amplitude map (top left), peak latency map (top right), reconstructed branches (bottom left), and velocity fits (bottom right) of the “MEA1k neuron” shown in blue in panel A. **C)** Axonal arbors on a DualMode device. **D)** Amplitude map (top left), peak latency map (top right), reconstructed branches (bottom left), and velocity fits (bottom right) of the “DualMode neuron” shown in blue in panel C.

In the DualMode recording, we found 58 units after spike-sorting and curation. Out of these, 51 had detectable axonal branches (shown in Figure 6C), and a total of 191 branches have been found (velocities: 368.88± 203.33 *mm/s*, path lengths: 504.18± 317.15 µm, *R*^2^ values: 0.95± 0.05). Similar to Figure 6B for the MEA1k neuron, Figure 6D shows detailed plots for one representative neuron displayed in blue in Figure 6C. For this unit, the channel selection yielded 2819 channels, where 14 axonal branches were traced featuring path lengths of 627.8± 426.24 µm, AP propagation velocities of 448.53± 173.33 *mm/s*, peak-to-peak extracellular amplitude of 103.2 µV, and *R*^2^ values of 0.96± 0.03.

We showed that the application of the proposed axonal reconstruction algorithm to spike-sorted data of HD-MEAs yields a high-throughput detection and assessment of axonal properties. The algorithm can potentially provide valuable information on axonal properties under physiological and pathological conditions.

## 4 Discussion

In this article, we introduce a novel, fully automated algorithm for reconstruction and AP-propagation velocity estimation of axons using HD-MEAs. The algorithm uses an efficient graph-based approach to reconstruct multiple axonal branches from extracellular electrical potential recordings. After detailing the different steps of the method, we assessed its performance using biophysical simulations. Afterwards, we validated our approach with experimental data recorded from two different HD-MEA devices - MEA1k and DualMode. We successfully reconstructed over 400 axonal branches and estimated the corresponding AP propagation velocities in two recording datasets. The developed algorithm and method can be used with all commercially available CMOS-based HD-MEAs. Moreover, we provide an open-source Python package available on GitHub (https://github.com/alejoe91/axon_velocity) and on PyPi (https://pypi.org/project/axon-velocity/) to facilitate the adoption of the method.

### Comparison with previous work

The presented algorithm builds upon previous approaches in our group to automatically reconstruct axonal arbors from HD-MEA extracellular signals. In *Yuan et al. 2020* [20], the authors introduced an axon-reconstruction method developed for the DualMode device. A basic idea of this approach that we also utilized for the method presented here, is to start the axon reconstruction backwards, i.e., from electrodes featuring late signal peak occurrences, which are most likely at the end of the respective axonal branches. However, a main limitation of this approach is that the search for axonal paths is *local*, i.e., that, in each step, the algorithm selects the next channel in the path only based on local signal amplitudes under the condition that the signal peak occurrence is earlier. This local search can result in zig-zag paths, as the algorithm has no information on the global structure of the signal landscape. To overcome this limitation, in *Ronchi et al. 2020* [34] we introduced a very first version of a graph-based algorithm. Subsequently, we made several improvements that were facilitated by the model-based validation that we present here. First, we extended the list of available filters for channel selection; in [34] only detection and kurtosis filters were used; second, we changed the interrogation of the graph to find axonal branches from using only the distance criterion, i.e., shortest distance (which could result in shortcuts and undetected axonal segments) to using a combination of distance and amplitude (***h***_***edge***_) criteria with the A* method; third, we changed the strategy to avoid duplicates in the path: instead of looking for and removing duplicate paths *a-posteriori*, we here utilized the set of neighboring channels to existing paths to avoid finding duplicates *a-priori*, which also resulted in a more efficient implementation. Finally, we added pruning, merging, and splitting steps that were not implemented in [34], which arguably provide a better estimation of the axon branches.

### Limitations

While the proposed method is, to the best of our knowledge, the first attempt of axonal tracing using HD-MEA signals in a fully-automated way and at high throughput, some limitations remain. Given the two-dimensional geometry of the recording electrode array, the method can only capture features in 2D and ignores modulations in the third dimension. A modulation in the z-distance of an axon to the MEA surface will result in a distorted estimate of axonal AP propagation velocity, as the distance traveled by the AP along a path in 3D will be different from its 2D projection onto the electrode plane. However, most neuronal preparations *in vitro* are 2D, at least most primary neuronal and organotypic cultures, where neurons and their neurites extend across a planar electrode array. Moreover, estimating the z-coordinate (the height above the electrode plane) in addition to the x-y coordinates of an axon is a complicated inverse problem. While the amplitude of the recorded axonal signal is known to depend on the position relative to the recording electrode, various other biophysical factors, such as ion-channel densities and kinetics, membrane capacitances, axial resistances, and axon geometries, can influence axonal AP conduction velocities in unmyelinated axons [43, 44, 45, 46, 47, 48]. In order to use the signal amplitude to correct for z-modulation, one would need to make assumptions about these other biophysical factors.

### Applications to neurological disease characterization and network dynamics

An automated and sufficiently accurate method to estimate axonal AP propagation velocities from HD-MEA recordings holds great promise to study axonal electrophysiology and pathophysiological conditions related to axonal dysfunction. A panoply of pathological conditions impair axonal functions and mostly result in conduction delays, which ultimately may cause conduction failures [49, 50, 51, 52, 53, 54].

Axonal dysfunction due to demyelination (e.g., multiple sclerosis) [55, 56], acute axonal damage [57], and channelopathies, among others, are shown to change axonal AP conduction properties [58, 59, 60]. Axonal features, such as differences in axon growth, axon signal conduction, time-course of axon degeneration or axon excitability can also be included in electrophysiological phenotypic characterization of human induced pluripotent stem cell (hiPSC)-derived neuronal cultures. Such cultures are available from patients suffering from neurological disorders and from healthy donors, so that electrophysiological biomarkers associated to neurological diseases can be established. In *Ronchi et al*. [34],for example, we made a first attempt to characterize axonal velocities of hiPSC-derived neuronal cultures and found significant differences between healthy motor and dopaminergic neurons and disease phenotypes featuring mutations related to Amyotrophic Lateral Sclerosis (ALS) and Parkinson’s disease (PD).

Besides identification and characterization of neurological diseases, an accurate determination of axonal AP propagation velocity opens up pathways to investigate axonal conduction times and delays and their role in neuronal coding and plasticity. Repetitive activity can alter the excitability of axonal membranes and AP conduction velocity, which can result in substantial changes in AP timings and spike propagation to presynaptic sites [61, 2, 62]. Conduction delays, which depend on conduction velocity and axonal length, can vary during repetitive activity, resulting in altered spike timings and intervals. Such changes in temporal spike patterns may be an important feature in shaping the neural code [63, 64, 65]. Similarly, axonal conduction velocities are highly adaptive in neuronal circuits and undergo changes in unmyelinated axons upon depolarization or during formation of new myelin sheaths depending on neuronal activity [66, 67, 68]. Our algorithm helps to facilitate the study of axonal conduction and potential failures, as it enables to simultaneously track a larger number of different axonal branches, to assess AP propagation velocities and conduction delays and to study the role of plasticity of conduction velocity in network-level dynamics. Such applications can also be extended to various model preparations such as organotypic cultures, acute brain slices and retinal slices.

### Outlook

In conclusion, in this article, we introduced and validated a novel automated method for axonal reconstruction from HD-MEA recordings, which enables to track changes in axonal conduction velocity over days. By providing an open-source Python package to use and apply the algorithm, we envision rapid adoption by the electrophysiology and HD-MEA community, which will eventually boost our understanding of biophysical and computational properties of axons in healthy and diseased states.

## 5 Acknowledgments

This work was supported by the ETH Zurich Postdoctoral Fellowship 19-2 FEL-17 (APB) and the ERC Advanced Grant 694829 “neuroXscales”.

## APPENDIX

### A Raw axonal branches estimation algorithm

In this appendix, we report the pseudo-code for the algorithm to estimate raw axonal branches from the constructed graph (see Section 2.2.3). Note that the pruning and merging steps are not included.

#### Algorithm 1: Identification of raw axonal paths from the graph.

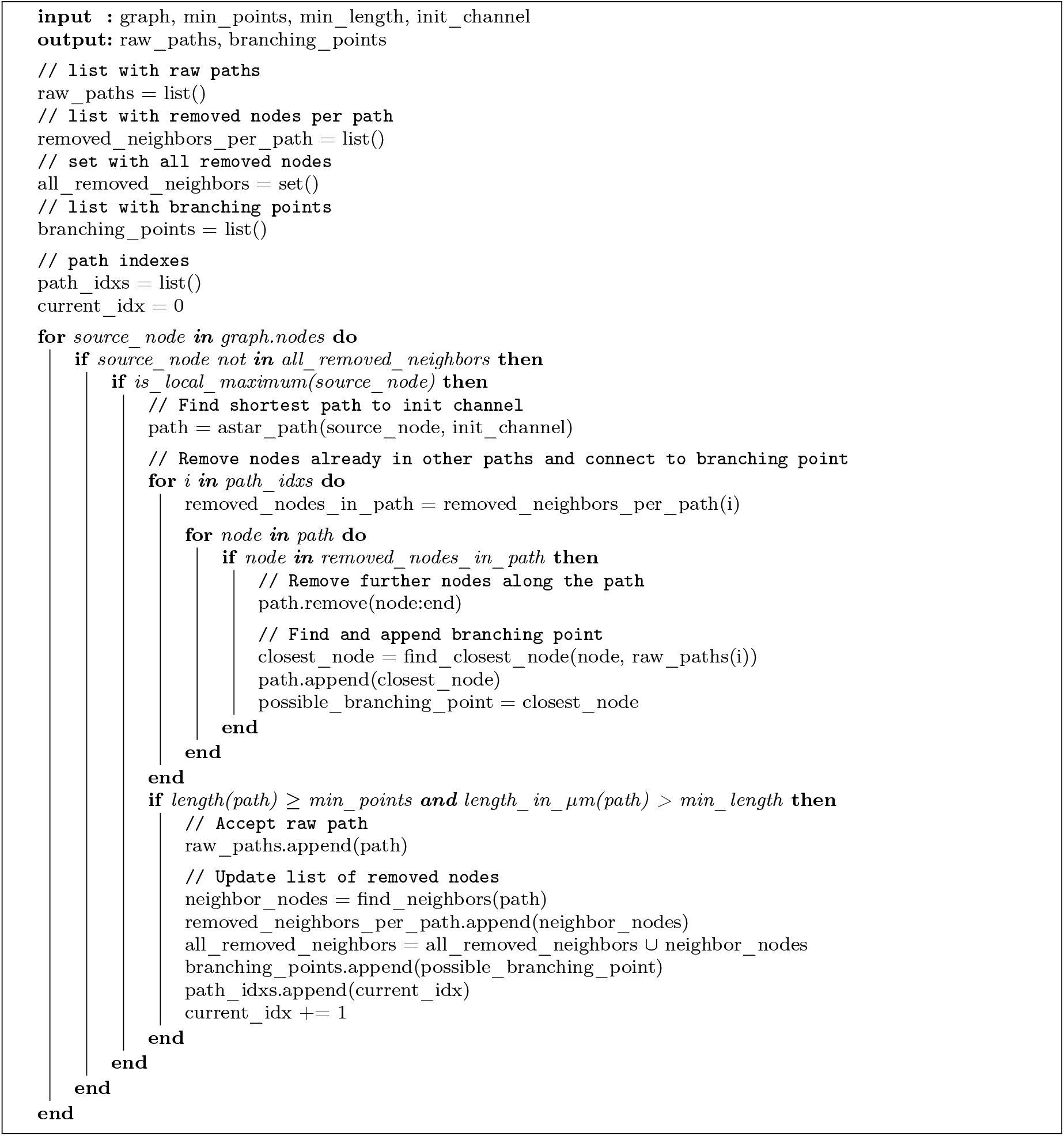

### B Description of parameters

In this appendix, we report a complete list of the parameters available for `axon_velocity` version The parameters are listed in Table 2.

**Table 2:**
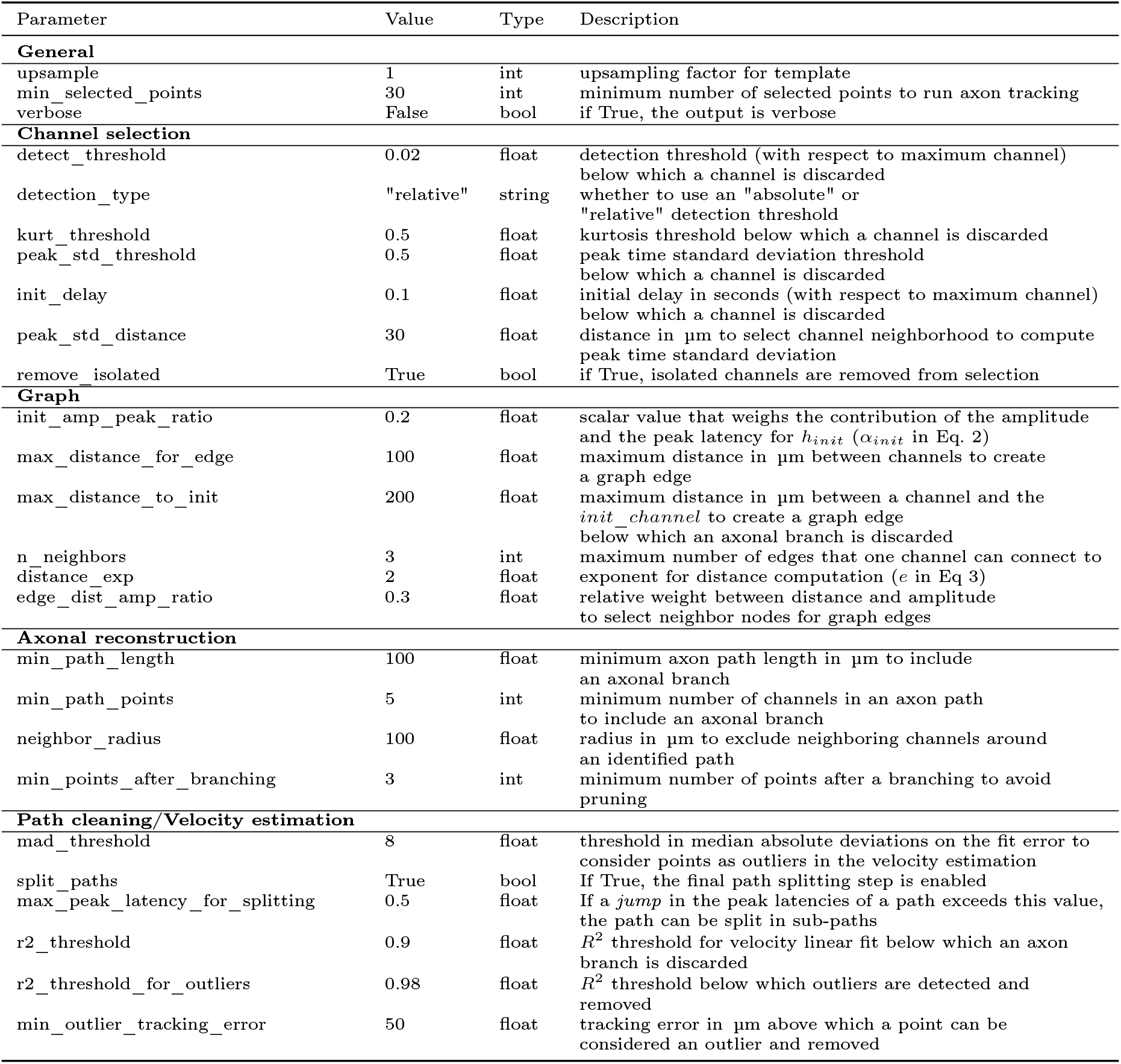
Additional parameters list for the `compute_graph_propagation_velocity()` function, including default values, data types, and descriptions.

## References

[1] A. L. Hodgkin and A. F. Huxley, “The components of membrane conductance in the giant axon of loligo,” The Journal of physiology, vol. 116, no. 4, p. 473, 1952.

[2] D. Debanne, “Information processing in the axon,” Nature Reviews Neuroscience, vol. 5, no. 4, pp. 304–316, 2004.

[3] K. S. Rockland, “What we can learn from the complex architecture of single axons,” Brain Structure and Function, pp. 1–21, 2020.

[4] T. Sasaki, N. Matsuki, and Y. Ikegaya, “Action-potential modulation during axonal conduction,” Science, vol. 331, no. 6017, pp. 599–601, 2011.

[5] T. van Kerkoerle, S. A. Marik, S. M. zum Alten Borgloh, and C. D. Gilbert, “Axonal plasticity associated with perceptual learning in adult macaque primary visual cortex,” Proceedings of the National Academy of Sciences, vol. 115, no. 41, pp. 10464–10469, 2018.

[6] Y. Shu, A. Hasenstaub, A. Duque, Y. Yu, and D. A. McCormick, “Modulation of intracortical synaptic potentials by presynaptic somatic membrane potential,” Nature, vol. 441, no. 7094, pp. 761–765, 2006.

[7] T. Sasaki, “The axon as a unique computational unit in neurons,” Neuroscience research, vol. 75, no. 2, pp. 83–88, 2013.

[8] J. A. Harrill, T. M. Freudenrich, D. W. Machacek, S. L. Stice, and W. R. Mundy, “Quantitative assessment of neurite outgrowth in human embryonic stem cell-derived hn2™ cells using automated high-content image analysis,” Neurotoxicology, vol. 31, no. 3, pp. 277–290, 2010.

[9] V. Laketa, J. C. Simpson, S. Bechtel, S. Wiemann, and R. Pepperkok, “High-content microscopy identifies new neurite outgrowth regulators,” Molecular biology of the cell, vol. 18, no. 1, pp. 242– 252, 2007.

[10] K.-M. Kim, S.-Y. Kim, J. Minxha, and G. T. R. Palmore, “A novel method for analyzing images of live nerve cells,” Journal of neuroscience methods, vol. 201, no. 1, pp. 98–105, 2011.

[11] A. Ossinger, A. Bajic, S. Pan, B. Andersson, P. Ranefall, N. Hailer, and N. Schizas, “A rapid and accurate method to quantify neurite outgrowth from cell and tissue cultures: Two image analytic approaches using adaptive thresholds or machine learning,” Journal of Neuroscience Methods, vol. 331, p. 108522, 2020.

[12] D. Wang, R. Lagerstrom, C. Sun, L. Bishof, P. Valotton, and M. Götte, “Hca-vision: Automated neurite outgrowth analysis,” Journal of biomolecular screening, vol. 15, no. 9, pp. 1165–1170, 2010.

[13] D. R. Hochbaum, Y. Zhao, S. L. Farhi, N. Klapoetke, C. A. Werley, V. Kapoor, P. Zou, J. M. Kralj, D. Maclaurin, N. Smedemark-Margulies, et al., “All-optical electrophysiology in mammalian neurons using engineered microbial rhodopsins,” Nature methods, vol. 11, no. 8, pp. 825–833, 2014.

[14] N. Ji, J. Freeman, and S. L. Smith, “Technologies for imaging neural activity in large volumes,” Nature neuroscience, vol. 19, no. 9, p. 1154, 2016.

[15] P. P. Laissue, R. A. Alghamdi, P. Tomancak, E. G. Reynaud, and H. Shroff, “Assessing phototoxicity in live fluorescence imaging,” Nature methods, vol. 14, no. 7, pp. 657–661, 2017.

[16] V. Emmenegger, M. E. J. Obien, F. Franke, and A. Hierlemann, “Technologies to study action potential propagation with a focus on hd-meas,” Frontiers in cellular neuroscience, vol. 13, p. 159, 2019.

[17] D. J. Bakkum, U. Frey, M. Radivojevic, T. L. Russell, J. Müller, M. Fiscella, H. Takahashi, and A. Hierlemann, “Tracking axonal action potential propagation on a high-density microelectrode array across hundreds of sites,” Nature communications, vol. 4, no. 1, pp. 1–12, 2013.

[18] T. Bullmann, M. Radivojevic, S. Huber, K. Deligkaris, A. R. Hierlemann, and U. Frey, “Largescale mapping of axonal arbors using high-density microelectrode arrays,” Frontiers in cellular neuroscience, vol. 13, p. 404, 2019.

[19] M. Radivojevic, F. Franke, M. Altermatt, J. Müller, A. Hierlemann, and D. J. Bakkum, “Tracking individual action potentials throughout mammalian axonal arbors,” Elife, vol. 6, p. e30198, 2017.

[20] X. Yuan, M. Schröter, M. E. J. Obien, M. Fiscella, W. Gong, T. Kikuchi, A. Odawara, S. Noji, I. Suzuki, J. Takahashi, A. Hierlemann, and U. Frey, “Versatile live-cell activity analysis platform for characterization of neuronal dynamics at single-cell and network level,” Nature Communications, vol. 11, no. 1, p. 4854, 2020.

[21] S. Ronchi, M. Fiscella, C. Marchetti, V. Viswam, J. Müller, U. Frey, and A. Hierlemann, “Singlecell electrical stimulation using cmos-based high-density microelectrode arrays,” Frontiers in neuroscience, vol. 13, p. 208, 2019.

[22] H. Lindén, E. Hagen, S. Leski, et al., “LFPy: a tool for biophysical simulation of extracellular potentials generated by detailed model neurons,” Frontiers in Neuroinformatics, vol. 7, p. 41, 2014.

[23] E. Hagen, S. Næss, T. V. Ness, and G. T. Einevoll, “Multimodal modeling of neural network activity: Computing lfp, ecog, eeg, and meg signals with lfpy 2.0,” Frontiers in neuroinformatics, vol. 12, 2018.

[24] N. T. Carnevale and M. L. Hines, The NEURON book. Cambridge University Press, 2006.

[25] N. W. Gouwens et al., “Systematic generation of biophysically detailed models for diverse cortical neuron types,” Nature communications, vol. 9, no. 1, p. 710, 2018.

[26] G. A. Ascoli, D. E. Donohue, and M. Halavi, “Neuromorpho. org: a central resource for neuronal morphologies,” Journal of Neuroscience, vol. 27, no. 35, pp. 9247–9251, 2007.

[27] S. Hallermann, C. P. De Kock, G. J. Stuart, and M. H. Kole, “State and location dependence of action potential metabolic cost in cortical pyramidal neurons,” Nature neuroscience, vol. 15, no. 7, pp. 1007–1014, 2012.

[28] T. V. Ness, C. Chintaluri, J. Potworowski, S. Leski, H. Głąbska, D. K. Wójcik, and G. T. Einevoll, “Modelling and analysis of electrical potentials recorded in microelectrode arrays (meas),” Neuroinformatics, vol. 13, no. 4, pp. 403–426, 2015.

[29] A. P. Buccino, M. Kuchta, K. H. Jæger, T. V. Ness, P. Berthet, K. A. Mardal, G. Cauwenberghs, and A. Tveito, “How does the presence of neural probes affect extracellular potentials?,” Journal of neural engineering, 2019.

[30] A. P. Buccino and G. T. Einevoll, “Mearec: a fast and customizable testbench simulator for ground-truth extracellular spiking activity,” Neuroinformatics, pp. 1–20, 2020.

[31] U. Frey, J. Sedivy, F. Heer, R. Pedron, M. Ballini, J. Mueller, D. Bakkum, S. Hafizovic, F. D. Faraci, F. Greve, et al., “Switch-matrix-based high-density microelectrode array in cmos technology,” IEEE Journal of Solid-State Circuits, vol. 45, no. 2, pp. 467–482, 2010.

[32] J. Müller, M. Ballini, P. Livi, Y. Chen, M. Radivojevic, A. Shadmani, V. Viswam, I. L. Jones, M. Fiscella, R. Diggelmann, et al., “High-resolution cmos mea platform to study neurons at subcellular, cellular, and network levels,” Lab on a Chip, vol. 15, no. 13, pp. 2767–2780, 2015.

[33] X. Yuan, V. Emmenegger, M. E. J. Obien, A. Hierlemann, and U. Frey, “Dual-mode microelectrode array featuring 20k electrodes and high snr for extracellular recording of neural networks,” in 2018 IEEE Biomedical Circuits and Systems Conference (BioCAS), pp. 1–4, IEEE, 2018.

[34] S. Ronchi, A. P. Buccino, G. Prack, S. S. Kumar, M. Schröter, M. Fiscella, and A. Hierlemann, “Electrophysiological phenotype characterization of human ipsc-derived neuronal cell lines by means of high-density microelectrode arrays,” Advanced Biology, p. 2000223, 2020.

[35] A. Hagberg, P. Swart, and D. S Chult, “Exploring network structure, dynamics, and function using networkx,” tech. rep., Los Alamos National Lab.(LANL), Los Alamos, NM (United States), 2008.

[36] F. Pedregosa, G. Varoquaux, A. Gramfort, V. Michel, B. Thirion, O. Grisel, M. Blondel, P. Prettenhofer, R. Weiss, V. Dubourg, J. Vanderplas, A. Passos, D. Cournapeau, M. Brucher, M. Perrot, and E. Duchesnay, “Scikit-learn: Machine learning in Python,” Journal of Machine Learning Research, vol. 12, pp. 2825–2830, 2011.

[37] X. Yuan, A. Hierlemann, and U. Frey, “Extracellular recording of entire neural networks using a dual-mode microelectrode array with 19,584 electrodes and high snr,” IEEE Journal of Solid-State Circuits, 2021.

[38] A. P. Buccino, C. L. Hurwitz, S. Garcia, J. Magland, J. H. Siegle, R. Hurwitz, and M. H. Hennig, “Spikeinterface, a unified framework for spike sorting,” Elife, vol. 9, p. e61834, 2020.

[39] N. A. Steinmetz, C. Aydin, A. Lebedeva, M. Okun, M. Pachitariu, M. Bauza, M. Beau, J. Bhagat, C. Böhm, M. Broux, et al., “Neuropixels 2.0: A miniaturized high-density probe for stable, longterm brain recordings,” bioRxiv, 2020.

[40] D. N. Hill, S. B. Mehta, and D. Kleinfeld, “Quality metrics to accompany spike sorting of extracellular signals,” Journal of Neuroscience, vol. 31, no. 24, pp. 8699–8705, 2011.

[41] C. Rossant, S. N. Kadir, D. F. Goodman, J. Schulman, M. L. Hunter, A. B. Saleem, A. Grosmark, M. Belluscio, G. H. Denfield, A. S. Ecker, et al., “Spike sorting for large, dense electrode arrays,” Nature neuroscience, vol. 19, no. 4, p. 634, 2016.

[42] C. Rossant, S. Kadir, D. Goodman, M. Hunter, and K. Harris, Phy, 2014. https://github.com/cortex-lab/phy.

[43] A. Hodgkin, “A note on conduction velocity,” The Journal of physiology, vol. 125, no. 1, pp. 221– 224, 1954.

[44] Y. Manor, C. Koch, and I. Segev, “Effect of geometrical irregularities on propagation delay in axonal trees,” Biophysical Journal, vol. 60, no. 6, pp. 1424–1437, 1991.

[45] G. M. Shepherd and K. M. Harris, “Three-dimensional structure and composition of ca3? ca1 axons in rat hippocampal slices: implications for presynaptic connectivity and compartmentalization,” Journal of Neuroscience, vol. 18, no. 20, pp. 8300–8310, 1998.

[46] K. Ganguly, L. Kiss, and M.-m. Poo, “Enhancement of presynaptic neuronal excitability by correlated presynaptic and postsynaptic spiking,” Nature neuroscience, vol. 3, no. 10, pp. 1018–1026, 2000.

[47] R. D. Fields, “Myelination: an overlooked mechanism of synaptic plasticity?,” The Neuroscientist, vol. 11, no. 6, pp. 528–531, 2005.

[48] Q. Cai, M. L. Davis, and Z.-H. Sheng, “Regulation of axonal mitochondrial transport and its impact on synaptic transmission,” Neuroscience research, vol. 70, no. 1, pp. 9–15, 2011.

[49] U. Suter and S. S. Scherer, “Disease mechanisms in inherited neuropathies,” Nature reviews neuroscience, vol. 4, no. 9, pp. 714–726, 2003.

[50] S. G. Waxman, “Axonal conduction and injury in multiple sclerosis: the role of sodium channels,” Nature Reviews Neuroscience, vol. 7, no. 12, pp. 932–941, 2006.

[51] C. Krarup and M. Moldovan, “Nerve conduction and excitability studies in peripheral nerve disorders,” Current opinion in neurology, vol. 22, no. 5, pp. 460–466, 2009.

[52] D. M. Kullmann, “Neurological channelopathies,” Annual review of neuroscience, vol. 33, pp. 151– 172, 2010.

[53] N. Egawa, J. Lok, K. Washida, and K. Arai, “Mechanisms of axonal damage and repair after central nervous system injury,” Translational stroke research, vol. 8, no. 1, pp. 14–21, 2017.

[54] S. Khalilpour, S. Latifi, G. Behnammanesh, A. M. S. A. Majid, A. S. A. Majid, and A. Tamayol, “Ischemic optic neuropathy as a model of neurodegenerative disorder: A review of pathogenic mechanism of axonal degeneration and the role of neuroprotection,” Journal of the neurological sciences, vol. 375, pp. 430–441, 2017.

[55] L. Steinman, “Multiple sclerosis: a coordinated immunological attack against myelin in the central nervous system,” Cell, vol. 85, no. 3, pp. 299–302, 1996.

[56] B. D. Trapp, R. Ransohoff, and R. Rudick, “Axonal pathology in multiple sclerosis: relationship to neurologic disability,” Current opinion in neurology, vol. 12, no. 3, pp. 295–302, 1999.

[57] D. H. Smith and D. F. Meaney, “Axonal damage in traumatic brain injury,” The neuroscientist, vol. 6, no. 6, pp. 483–495, 2000.

[58] P. J. Goadsby, P. R. Holland, M. Martins-Oliveira, J. Hoffmann, C. Schankin, and S. Akerman, “Pathophysiology of migraine: a disorder of sensory processing,” Physiological reviews, 2017.

[59] J. Oyrer, S. Maljevic, I. E. Scheffer, S. F. Berkovic, S. Petrou, and C. A. Reid, “Ion channels in genetic epilepsy: from genes and mechanisms to disease-targeted therapies,” Pharmacological reviews, vol. 70, no. 1, pp. 142–173, 2018.

[60] D. Pietrobon, “Ion channels in migraine disorders,” Current Opinion in Physiology, vol. 2, pp. 98– 108, 2018.

[61] J. R. Geiger and P. Jonas, “Dynamic control of presynaptic ca2+ inflow by fast-inactivating k+ channels in hippocampal mossy fiber boutons,” Neuron, vol. 28, no. 3, pp. 927–939, 2000.

[62] S. Boudkkazi, L. Fronzaroli-Molinieres, and D. Debanne, “Presynaptic action potential waveform determines cortical synaptic latency,” The Journal of physiology, vol. 589, no. 5, pp. 1117–1131, 2011.

[63] E. M. Izhikevich, “Polychronization: computation with spikes,” Neural computation, vol. 18, no. 2, pp. 245–282, 2006.

[64] D. Bucher and J.-M. Goaillard, “Beyond faithful conduction: short-term dynamics, neuromodulation, and long-term regulation of spike propagation in the axon,” Progress in neurobiology, vol. 94, no. 4, pp. 307–346, 2011.

[65] D. Bucher, “Contribution of axons to short-term dynamics of neuronal communication,” in Axons and Brain Architecture, pp. 245–263, Elsevier, 2016.

[66] D. Debanne, E. Campanac, A. Bialowas, E. Carlier, and G. Alcaraz, “Axon physiology,” Physiological reviews, vol. 91, no. 2, pp. 555–602, 2011.

[67] R. De Col, K. Messlinger, and R. W. Carr, “Repetitive activity slows axonal conduction velocity and concomitantly increases mechanical activation threshold in single axons of the rat cranial dura,” The Journal of physiology, vol. 590, no. 4, pp. 725–736, 2012.

[68] I. A. McKenzie, D. Ohayon, H. Li, J. P. De Faria, B. Emery, K. Tohyama, and W. D. Richardson, “Motor skill learning requires active central myelination,” science, vol. 346, no. 6207, pp. 318–322, 2014.

